# Prediction of Ca^2+^ binding site in proteins with a fast and accurate method based on statistical mechanics and analysis of crystal structures

**DOI:** 10.1101/2024.03.03.583063

**Authors:** Abdul Basit, Devapriya Choudhury, Pradipta Bandyopadhyay

## Abstract

Predicting the precise locations of metal binding sites within metalloproteins is a crucial challenge in biophysics. A fast, accurate, and interpretable computational prediction method can complement the experimental studies. With this endeavor, In the current work, we have developed a method to predict the location of Ca^2+^ ions in calcium-binding proteins using a physics-based method with an all-atom description of the proteins, which is substantially faster than the molecular dynamics simulation-based methods with accuracy as good as data-driven approaches. Our methodology uses the three-dimensional reference interaction site model (3D-RISM), a statistical mechanical theory, to calculate Ca^2+^ ion density around protein structures, and the locations of the Ca^2+^ ions are obtained from the density. We have taken previously used datasets to assess the efficacy of our method as compared to previous works. Our accuracy is found to be 88%, comparable with the FEATURE program, one of the well-known data-driven methods. Moreover, our method being physical, the reasons for failures can be ascertained in most cases. We have thoroughly examined the failed cases using different structural and crystallographic measures, such as B-factor, R-factor, electron density map, and geometry at the binding site. It has been found that X-ray structures have issues in many of the failed cases, such as geometric irregularities and dubious assignment of ion positions. Our algorithm, along with the checks for structural accuracy, is a major step in predicting calcium ion positions in metalloproteins.

## 1. Introduction

Metalloproteins are ubiquitous in biology and involve a large number of functions, such as electron transfer, oxygen transport, enzymatic catalysis, DNA replication and repair, and signal transduction. One of the outstanding challenges in structural biology is the determination and prediction of the positions of the metal ions in proteins. Although there are thousands of metalloproteins in the PDB, experimental determinations and computational predictions face several issues, and continuous improvement of both is needed. As the coordination geometry of metal ions varies widely, from simple to complex arrangements, making the exact geometry prediction a non-trivial task. Metal binding sites can be dynamic, responding to cellular signals and inducing conformational changes, requiring dynamic studies for a comprehensive understanding ^1^. There can be multiple challenges in X-ray crystallography regarding the determination of the positions of the ions. For instance, the process of purification and crystallization can be particularly problematic for some proteins. During these processes, non-native metals may substitute native ions, and buffer components and crystallization agents can influence metal binding ^2^. Crystal packing forces can perturb metal ion positions, and crystal quality can lead to lower-quality diffraction data ^3^. Sometimes, the metal’s redox state is altered, which can lead to a change in coordination geometry ^4^. X-ray radiation can cause damage to crystals during data collection, potentially displacing or dislodging metal ions ^5-6^. Solvent molecules, like water, can also coordinate with metal ions, posing challenges in distinguishing between water and protein ligands ^7^. On the other hand, for nuclear magnetic resonance (NMR) investigations, analyzing metal ions binding to proteins involves examining the chemical shift perturbations induced by the interaction with metal ions. Diamagnetic metal ions, being NMR silent, do not provide insights into the geometric structure of the metal-binding sites. Paramagnetic metal ions within proteins, characterized by unpaired electrons, influence protein nuclei’s chemical shifts and relaxation rates. The metal may affect signals from nuclei in proximity to the metal ion even beyond detection, making it challenging to detect information about the geometric structure of the protein near the metal site. The presence of the metal ion can influence signals from nuclei in proximity, even beyond detection, thereby complicating the extraction of information regarding the geometric structure of the metal site ^8-9^. Deciphering paramagnetic NMR data poses challenges due to its complexity. Detailed interpretation often demands extensive and versatile datasets. Consequently, paramagnetic NMR data are frequently utilized semi-quantitatively, relying on reduced datasets and approximations for interpretation ^10-11^.

An accurate computational predictive method can complement experimental results and, in some cases, be useful when experimental structures are unavailable. Most of the current prediction methods are data-driven, which hinge on the extraction of features from both amino acid sequences and structures to facilitate the training of predictive models ^12^. Some approaches centred around sequences ^13-14^, and others exhibit a structural emphasis akin to methods like Fold-X ^15^, MIB ^16-17^, and BioMetAll ^18^. Other methodologies take a dual approach, combining sequence-based and structural information to achieve a more comprehensive analytical standpoint ^19-20^. AlphaFold predicts protein 3D structures, focusing on the spatial arrangement of amino acids in the holo conformations ^21^. However, It does not provide explicit predictions for the coordinates of ligands, cofactors, metals, or water molecules associated with the protein structures. AlphaFill utilizes homology to overcome this limitation and transplants missing exogenous ligands from similar PDBs, guided by a sequence identity cutoff of 25% ^22^. RoseTTAFold is being made general to predict structures of proteins with small molecules and metal ions ^23^.

Among different metals, we have chosen Ca^2+^ ion as our metal of interest in the current work. Ca^2+^ is one of the most important metal ions in biology as it acts as a secondary messenger in several regulatory processes, such as muscle contraction, neurotransmission, release, gene transcription, and cell signalling cascades ^24^. Calcium ion’s unique and special characteristics include high charge density, exclusive coordination by oxygens, and high specificity ^25^. Additionally, their role in cell signalling pathways and the ability to induce conformational changes in proteins, primarily through dynamic interactions, contribute to their distinct significance ^26^. Predicting the calcium-binding site within a protein with a weak affinity for Ca^2+^ can pose challenges for X-ray crystallography ^27-28^. Phasing difficulties may arise when calcium ions are the sole anomalous scatterers ^29^. While existing literature demonstrates single-wavelength anomalous diffraction (SAD) phasing with calcium ions, the process is anticipated to be challenging ^30^. The challenge in NMR spectroscopy of calcium-binding proteins arises from the low natural abundance and resonance frequency of the only spin-active calcium isotope, ^43^Ca ^31^. Another challenge in identifying Ca^2+^ binding sites stems from the fact that the positions of the oxygen atoms, which chelate the Ca^2+^, are frequently not directly established. This limitation arises from the isotopically abundant ^16^O, which inherently possesses a zero nuclear spin ^32^.

Among the computational works for predicting the positions of Ca^2+^ ions, Jenny Yang’s group has developed a calcium-binding site prediction algorithm rooted in graph theory ^33–37^. The FEATURE program-based algorithm from Altman’s group, requiring physiochemical descriptors, has also been leveraged for calcium-binding site predictions ^38–42^. Mazumder et al. employed Support Vector Machines (SVM) to classify binding sites into EF and non-EF categories and predict the affinity and design based on their sequence pattern ^43-44^. For in-depth information on these methods and their applications, readers are encouraged to refer to related review articles for a comprehensive overview of the evolving field of metal-binding site prediction in proteins ^45–50^. There are also physics-based approaches that rely on physical models, such as molecular dynamics (MD) simulations ^51^ and Poisson-Boltzmann electrostatics ^52^. MD simulation finds application in predicting binding sites for metal ions, including calcium, but it can demand substantial computational resources and time ^53^ and is not suitable as a prediction tool for a database. Moreover, getting convergence in MD simulations with ions can be difficult ^54^.

In the current work, we have developed a computational protocol that circumvents the problems associated with physics-based predictions with accuracy as good as the best data-driven methods. This method is (a) orders of magnitude faster than MD-based approaches, gives accuracy as good as the FEATURE software when applied to databases used previously for predicting Ca^2+^ positions, (c) the method being physical, reasons for success and failure can be ascertained in most cases which can be difficult for data-driven methods. Our method uses the 3D-reference interaction site model (3D-RISM), a statistical mechanical theory on energy-minimized structures of the proteins. 3D-RISM has been used previously for predicting the location of ions in a protein ^55-60^; however, as far as our knowledge goes, this is the first time this method has been used for predicting ion locations in a large-scale study. To probe deeper into the success and failures of our technique, we have checked the quality of the protein crystal structures present in the databases considered in this work by R-factor, B-factor and electron density maps. We achieved an accuracy of 81% when the cutoff distance of 2 Å was used between the position of the Ca^2+^ in the protein structure and that obtained from the prediction model. The accuracy increases to 88% if the cutoff is made 3.5 Å. We dissected the reasons for the cases where the algorithm failed and found that, in most cases, inaccuracy in the X-ray structures is the main reason for it.

The manuscript follows this structure. The next section gives the methodology of our work, followed by results and discussion. It concludes with a summary and avenues for future work.

## 2. Methodology

### 2.1 3D-RISM method

In statistical mechanics, the pair-correlation function (PCF) between two particles originates from the two-body reduced distribution function. It is defined as the ratio of the probability of finding two particles at a certain separation (for non-spherical particles, orientation between the two particles is also needed for a complete description) to that in the bulk. In the current problem, a high value of PCF of Ca^2+^ around specific regions of the proteins indicates a higher probability of finding Ca^2+^ there than the concentration of the ions in the system (which is the bulk density).

Calculation of PCF is being done extensively with liquid state integral equation theories in studying fluid properties ^61^C. Calculating PCF facilitates the calculation of all thermodynamic properties if the inter-particle interactions are pairwise. In 1972, a new PCF-based theory for non-spherical systems was proposed by Chandler and Anderson and termed it as the reference interaction site model (RISM) ^62^. The details of the RISM method can be seen from previous literature ^63^. A brief introduction to the PCF theory for a one-component system is given in the following.

The PCF-based method solves the Ornstein-Zernike (OZ) equation with a closure ^61,64^. In a one-component spherical system with density *ρ* the OZ equation takes the following form

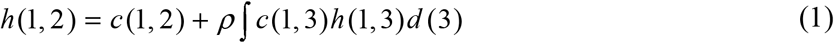

Where the PCF, *g* (1, 2) = *h*(1, 2) + 1

Where h and c represent total and direct correlation functions, respectively, and the numbers inside the parenthesis represent the coordinates. Equation (1) has two unknowns, necessitating an additional equation called closure to establish the relationship between h and c and to get the unknowns. The general closure relationship is expressed by

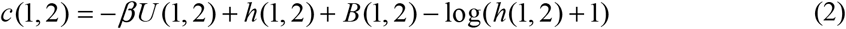

Where U represents the potential energy, β=1/kT (where k is the Boltzmann constant and T is absolute temperature), and B(1,2) are a set of expressions called bridge functions. As evaluating the bridge functions is challenging, different closures approximate it differently. The most straightforward closure, i.e., hypernetted chain (HNC), considers bridge terms zero.

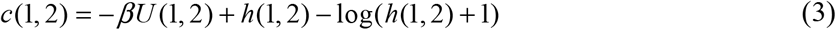

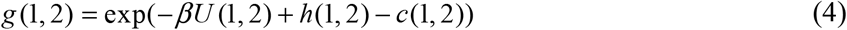

As the HNC closure finds convergence of the iterative procedure used in the PCF theory difficult, the Kovalenko-Hirata (KH) ^65^ closure is used in this work. KH closure can be defined as follows:

Equation (4) can be written as

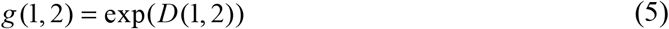

where

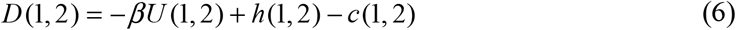

In Kovalenko-Hirata (KH) closure,

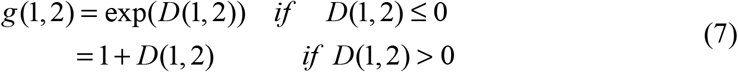

In RISM, a molecule is modelled by multiple sites and orientations are integrated out, so the PCF depend only on the inter-site distances.

In the 3D-RISM method, to get the solvent density distribution (including ions) around the solute over a grid, the solvent coordinates are averaged over fixed solute geometry ^65–67^. For the details of the 3D-RISM method, previous literature can be seen ^63^. The assignment of the density distribution function obtained from 3D-RISM to the population of a species (ions and waters) was done by the Placevent algorithm ^68^. The primary objective of this algorithm is to transform the 3D RISM distribution function into a population function that facilitates the representation of a discrete distribution of explicit solvent atoms. This is achieved by integrating the PCF over distance until the result of integration (i.e. population) becomes one. This integration starts with the highest value of the PCF (g_max_(**r**), **r** is the centre of the calcium ion), giving rise to the population of one Ca^2+^ ion and for other Ca^2+^ ions, these integrations are done for the subsequent highest value of the PCF (the region that gives the previous population is not considered to get the current highest value of the PCF) to generate the location of the other Ca^2+^ ions. The population of different calcium ions can be ranked based on the values of the PCF at their centres. We have used the Placevent algorithm to get the location of Ca^2+^ and water.

### 2.2 Evaluating the quality of X-ray structures of proteins

To evaluate the quality of a protein structure, a diverse array of methods and validation tools are at the disposal of researchers, each offering insights into the accuracy of the protein structure ^69^. The R-factors are quantitative indicators of the concordance between the model and experimental data, where lower values denote a more robust alignment ^70^. The Ramachandran plot is a cornerstone in model evaluation, which can graphically portray backbone dihedral angles. Ramachandran outliers refer to amino acids with unfavourable dihedral angles; the plot is a powerful tool for identifying them ^71^. Clash analysis involves scrutinizing the occurrence of steric clashes between atoms within the model. B-factor analysis delves into atomic displacement parameters, where lower values signify well-defined positions ^72^. Exploring electron density maps further elucidates the agreement between the model and empirical data, which can be complex and challenging ^73^. Validation software such as PROCHECK ^74^, MolProbity ^75^, and Coot ^76^ and servers like CheckMyBlob ^77^ and CheckMyMetal ^78^ facilitate the automated assessment of structural parameters. Moreover, the biological context and functional relevance are crucial when analyzing protein structures ^33,79-80^.

R-values: R-factor (or R-value) quantifies the discrepancy between the observed experimental data and the calculated intensities from diffractions. We estimate it using the formula:

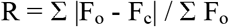

The equation involves the observed or experimental structure factor amplitude (F_o_), the calculated structure factor amplitude (F_c_) based on the atomic model, and a summation of all reflections of X-rays.

B-factor: The B-factor, also called the temperature factor or atomic displacement parameter, measures atoms’ thermal motion or disorder in a protein or other macromolecule. The equation represents the B-factor is:

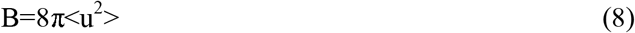

Where ‘B’ is the B-factor, and <u^2^> is the mean square displacement of the atom from its average position.

### 2.3 Datasets for Ca^2+^ Binding Site: Evaluation of Prediction and Structure Quality

To assess the effectiveness of our protocol, we compiled three datasets for testing. Dataset-A was prepared from previously used datasets ^42,81–83^. Dataset-B is a refined version of Dataset-A, obtained by applying structural quality filters to enhance its quality, described in the next section. Subsequently, we prepared Dataset C in-house with high-quality structures. Detailed characteristics of each dataset are as follows.

#### Dataset-A: from previous works

We assembled the initial dataset from four previously conducted studies. The first, from Zhou et al., encompassed 312 proteins ^42^; the second, from Nayal et al., comprised 63 protein structures ^81^; the third, compiled by Liang et al., included 40 PDB entries ^82^; the fourth, by Pidcock and Moore, consisted of 44 proteins ^83^. These studies had common protein structures, and we curated a non-redundant set of 134 single calcium-binding proteins for our analysis.

#### Dataset-B: filtered data

We prepared a refined dataset by applying quality filtration to the first dataset, i.e. dataset-A, to alleviate inherent artefacts in PDB structures. The filtration criteria included resolution values lower than 2 Å and R-factors below 0.20, ensuring their B-factors oxygens coordinated to calcium adhered to expected norms. The validation of electron density on calcium sites was an integral step in this filtration process. We also confirmed the chemical identity of the ion using the calcium bond-valence sum approach described later ^84^. Figure 1 illustrates the sequential steps of this data filtration procedure. These filtration steps culminated in a refined subset of nine proteins. We have also investigated the role of separate filters on prediction accuracy, the criteria of which are outlined in the results section.

**Figure 1:**
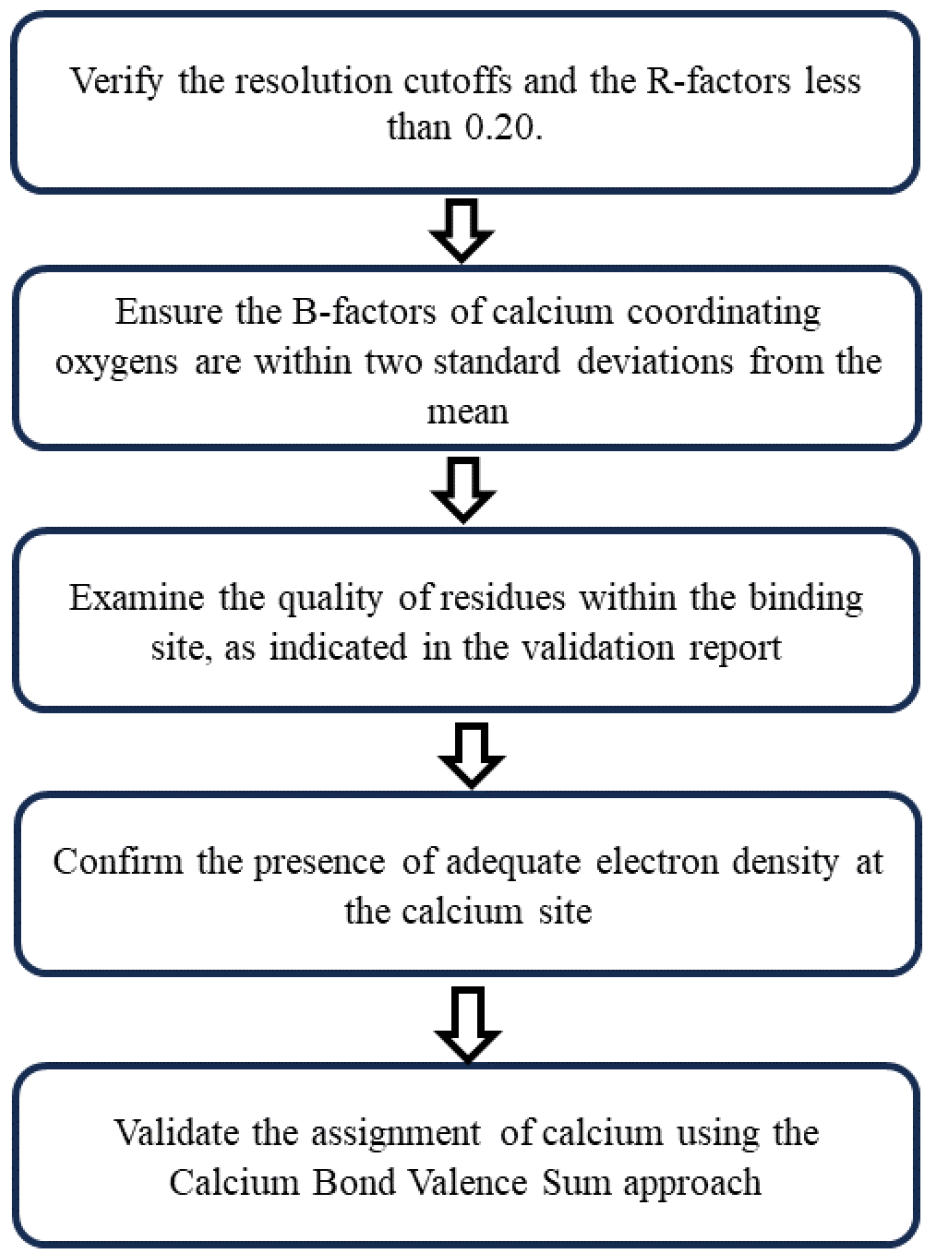
llustration of the steps involved in the filtration of protein structures. This process is a series of stages or actions to select and refine the protein structures for the binding site predictions. We designed each step in the filtration process to remove irrelevant or problematic proteins, guaranteeing that only the most suitable candidates are eligible for further examination.

#### Dataset-C: Dataset of High-Quality Structures (in-house prepared)

We executed a curation process to pursue an even higher-quality new dataset of calcium-binding protein structures. The initial phase encompassed a search within MetalPDB ^85^, focusing on PDB entries having Ca^2+^ ions. Afterwards, we initiated a filtration procedure that included a maximum sequence identity threshold of 70%, with criteria for resolution up to 1.25 Å ^86^. The other measures are the same as shown in Figure 1. We excluded Instances characterized by coordination to fewer than four ligands. We completed binding site geometries whenever required by utilizing biological assembly or symmetry mates (e.g., for the PDB id 4RU3). As a culmination of these iterative steps, we curated 33 single calcium-binding proteins.

For all the datasets, we considered only one for the subsequent calculations in the case of multiple identical calcium sites within a protein. In cases where there are multiple conformations for a residue within a protein structure, we utilize the first conformation. We chose these proteins when no exogenous ligand other than water was present in their active sites. For a detailed list of PDB IDs included in these datasets, please refer to Table 1 in the supporting information section.

**Table 1:** Impact of Structural Filters on Protein Structure Dataset: The table illustrates the influence of various structural filters, such as resolution, R-factors, B-factor, geometric irregularities, electron density, and CVBS, on the number of protein structures showcasing the resulting structure reduction after filtration.

| **Filters** | **Characteristic** | **Number of Structures** | **Success Rate(%)** |
| --- | --- | --- | --- |
| No filter | - | 134 | 85 |
| filter-1 | Resolution | 76 | 87 |
| filter-2 | R-factors | 33 | 94 |
| filter-3 | B-factor | 32 | 94 |
| filter-4 | Validation report | 19 | 84 |
| filter-5 | Electron density | 18 | 89 |
| filter-6 | CVBS | 9 | 100 |

### 2.4 The Calcium Bond Valence Sum (CBVS) Approach

The calcium bond-valence sum is a method to discern metal ion identity within protein structures ^84^. CBVS focuses on analyzing metal binding sites and evaluating the quality of metal-ion coordination based on bond valence principles. The bond valence sum involves adding up the valences of all the atoms directly bonded to a central metal ion. Before summing them up, calculated valences are multiplied by their occupancies (p_i_) in the crystal structure of the protein.

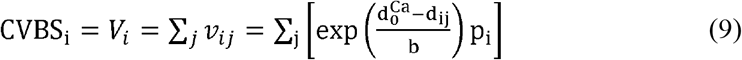

In the above expression, d_0_ represents the ideal bond length between Ca^2+^ and its ligand, and d_ij_ is the observed bond length of metal i and ligand j. Factor b is an empirical parameter with a value of 0.37 Å that accounts for the strength of interactions between the Ca^2+^ and its ligands. The calculated bond valence sum is then compared with the expected valence sum for Ca^2+,^ i.e. 2. If the calculated and expected values agree, it confirms the ion is Ca^2+^; otherwise, there are different metals according to CBVS scores.

### 2.5 Model and Parameters

Molecular Mechanics: In the simulations, we used the PMEMD module of the AMBER 20 package to perform the calculations. We used ff14SB forcefield for proteins ^87^. We implicitly considered the water around the proteins with Hawkins, Cramer, and Truhlar’s pairwise Generalized Born model and Tsui and Case parameters ^88^. Next, we conducted a minimization process for each protein using the steepest descent algorithm, completing 1000 minimization steps. We removed every heteroatom, including Ca^2+^ ions and water molecules, for this relaxation step.

1D-RISM: We employed the “rism1d” module from the AMBER 20 package to conduct a 1D-RISM calculation. The selected water model was SPC/E ^89^, and we used ion parameters based on Joung-Cheatham ^90^ and Li/Merz ^91^. We did all calculations at a temperature of 298 K. In this process, we considered the 0.1 M CaCl^2^ and adjusted the water density accordingly^92^.

3D-RISM: In the subsequent phase, we harnessed the solvent characteristics derived from the 1D-RISM computation to execute a 3D-RISM calculation utilizing a distinct module termed “rism3d.snglpnt” from the AMBER 20 package. During the 3D-RISM calculation, we implemented the Kovalenko−Hirata (KH) closure method. We prepared the simulation boxes, ensuring a minimum distance of 30.0 Å between any protein atom and the boundary or edge of the box. During all these calculations, grid spacing was 0.5 Å.

Placevent: The placevent algorithm needs the concentration of solvent species as input to convert them into explicit atoms.

## 3. Results

Our datasets unveiled a spectrum of coordination geometries from tetrahedral to octahedral within the chosen protein structures to predict calcium-binding sites. The binding site of each protein has a combination of various residues, contributing to a diversity of charges. The aspartic acids (ASP) and glutamic acid (GLU) are the most prevalent within these coordinating residues. Polar residues manifested a higher frequency within these binding sites than their hydrophobic counterparts. Moreover, many protein structures indicated the presence of water molecules snugly nestled within the binding sites. Nonetheless, it is noteworthy that several structures demonstrated a low coordination number (≤4) in the first dataset. Figures S1 and S2 show site Ca^2+^ coordinations, water within binding sites, and the distribution of amino acids.

Figure 3 presents the visualization of the Ca^2+^ density around three proteins obtained from 3D-RISM calculations using an isosurface representation of the PCF (taken as four) in VMD ^93^. The distributions were transformed into Ca^2+^ ions using the placevent algorithm and subsequently compared with the documented positions.

**Figure 2.**
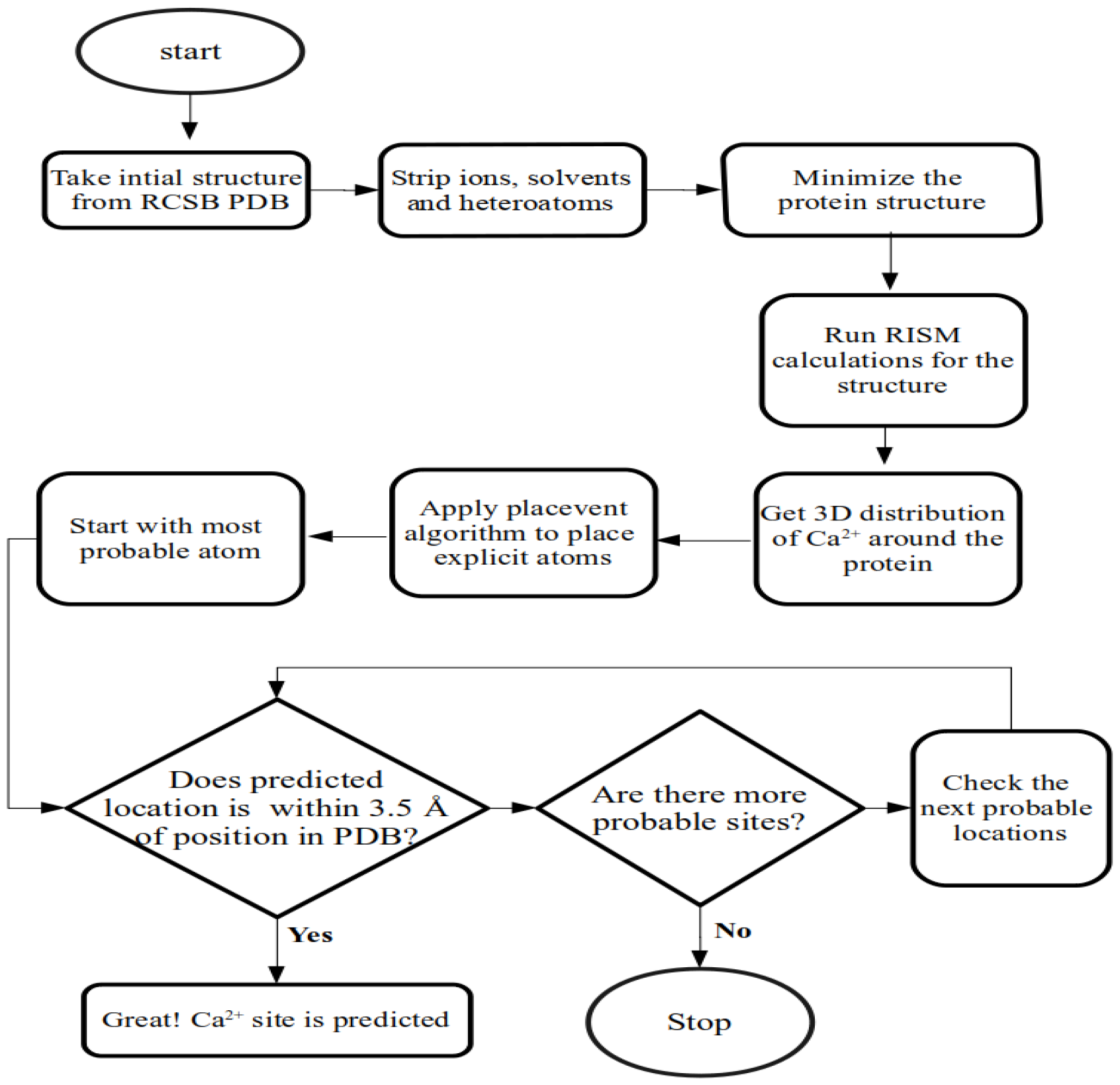
The figure illustrates the different steps involved in the calcium-binding site prediction algorithm.

**Figure 3:**
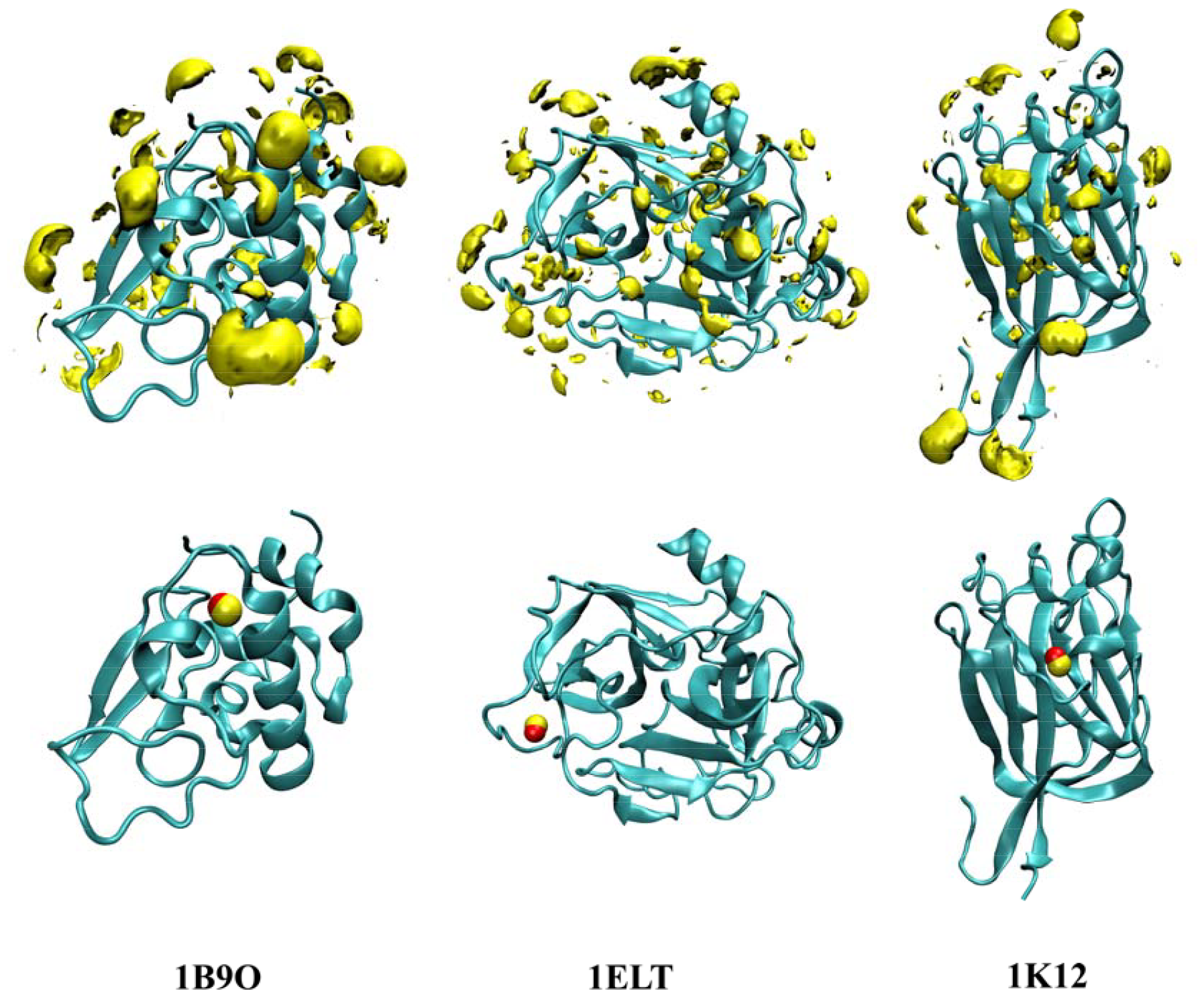
Representative outcome of our prediction algorithm for three different protein structures, identified by their PDB IDs, i.e. 1B9O, 1ELT and 1K12. The upper panel shows the distribution of Ca^2+^ ions around these proteins in yellow, obtained from 3D-RISM calculations. The lower panel shows the predicted and documented locations of Ca^2+^ ions around the same structures in yellow and red, respectively. All these three predicted Ca^2+^ are within 1 Å of the actual position.

### 3.1 Performance Evaluation of Prediction Algorithms: Insights from Metrics

We first assessed the prediction algorithms using recall (sensitivity). This evaluation entailed comparing the predicted positions of Ca^2+^ with documented sites obtained from experimental data. Recall signifies the proportion of true positive predictions among all positive cases. Figure 4 presents successful instances and corresponding success rates at different cutoffs for successful prediction. Increasing the cutoff value enhances the success rate of the algorithm. Notably, at a cutoff of 3.5 Å, the success rate is 88%. The subsequent sections will delve into the results, all of which are examined based on a 3.5 Å cutoff unless specified otherwise.

**Figure 4:**
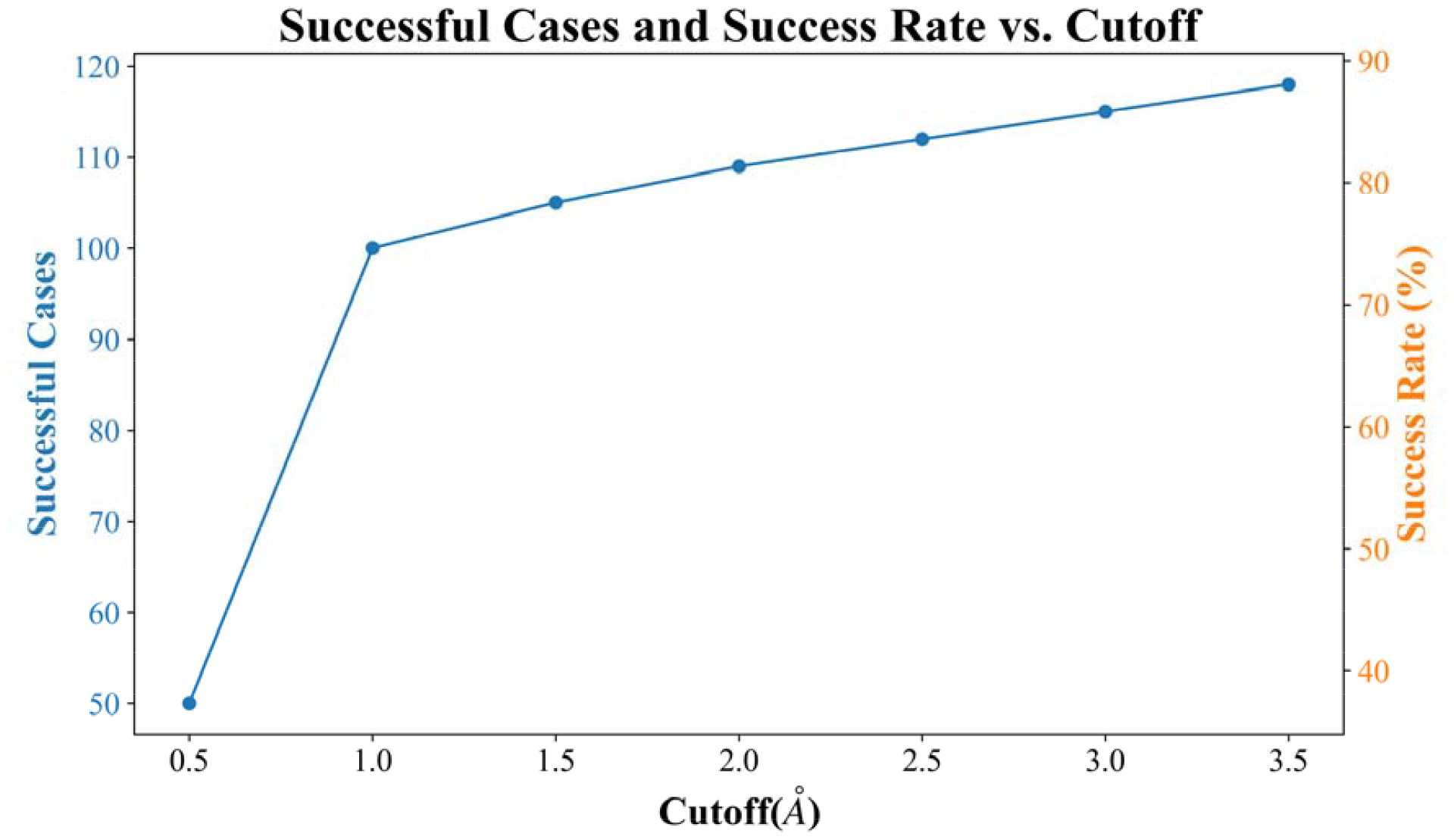
Relationship between cutoff values and successful cases and success rates. The line plot illustrates the variation in the number of successful cases and the corresponding success rates as a function of different cutoff values. As the cutoff value increases, the number of successful cases and success rates exhibit an upward trend.

#### Assessing Spatial Precision in Calcium-Binding Site Prediction

We also assessed the prediction algorithm using deviation in the distance between predicted and reported positions on the correctly identified location to quantify the precision. The average deviation is 0.76 Å, and the median deviation is 0.53 Å. It means most predicted positions are close to the calcium sites, with distances falling within 1 Å. The binding site prediction algorithm encounters challenges in cases where certain conditions are not met, such as the binding site being on the surface (e.g. for PDB id 1GSP), the absence of negatively charged residues or lower coordination of Ca^2+^ ions (e.g. for PDB id 1M1U). In some cases, we observed that the predicted binding site had equal or more negatively charged residues than the actual binding site (e.g., 1BJI). However, it is worth noting that the method successfully identifies calcium sites incorporating one or more water molecules, as seen in structures like 1J5U (Figure S3).

#### Evaluating Ca^2+^ Prediction: First Rank Percentage

The evaluation of predicted Ca^2+^ ion positions also involves the consideration of the first rank percentage (FRP) as a crucial metric. FRP refers to the percentage of instances where a structure achieves the highest rank prediction as the accurate prediction. The ranking of predicted Ca^2+^ ions based on their respective g(**r**) values forms the basis of our analysis. Our protocol demonstrated 84% success in correctly identifying and ranking calcium sites as the first predictions. However, it is noteworthy that for the successful predictions the first-ranked ion missed the binding sites of just five structures, where it was the second ion in three cases and the third in two.

### 3.2 Analyzing Unsuccessful Predictions

Out of the 134 cases considered, there were 16 instances in which the predictions were unsuccessful. Despite these unsuccessful predictions, it is essential to highlight that the binding sites still exhibit significant g(**r**) values, indicating the presence of potential interaction sites. Hence, these instances cannot be termed as complete failures. Figure 5 shows the PCF in the protein binding site, g_site_(**r**_**1**_) and maximum PCF values, i.e. g_max_(**r**_2_). In the following sections, we did further investigations to understand these specific cases better and develop enhanced methods for accurately predicting metal-binding sites in such challenging scenarios. It can be seen from the figure there are several instances where PCF at the binding site is close to the maximum PCF. For instance, for the PDB ID, 1KUH, these values are very close. However, the average distance between **r**_**1**_ and **r**_**2**_ is about 24 Angstrom.

**Figure 5:**
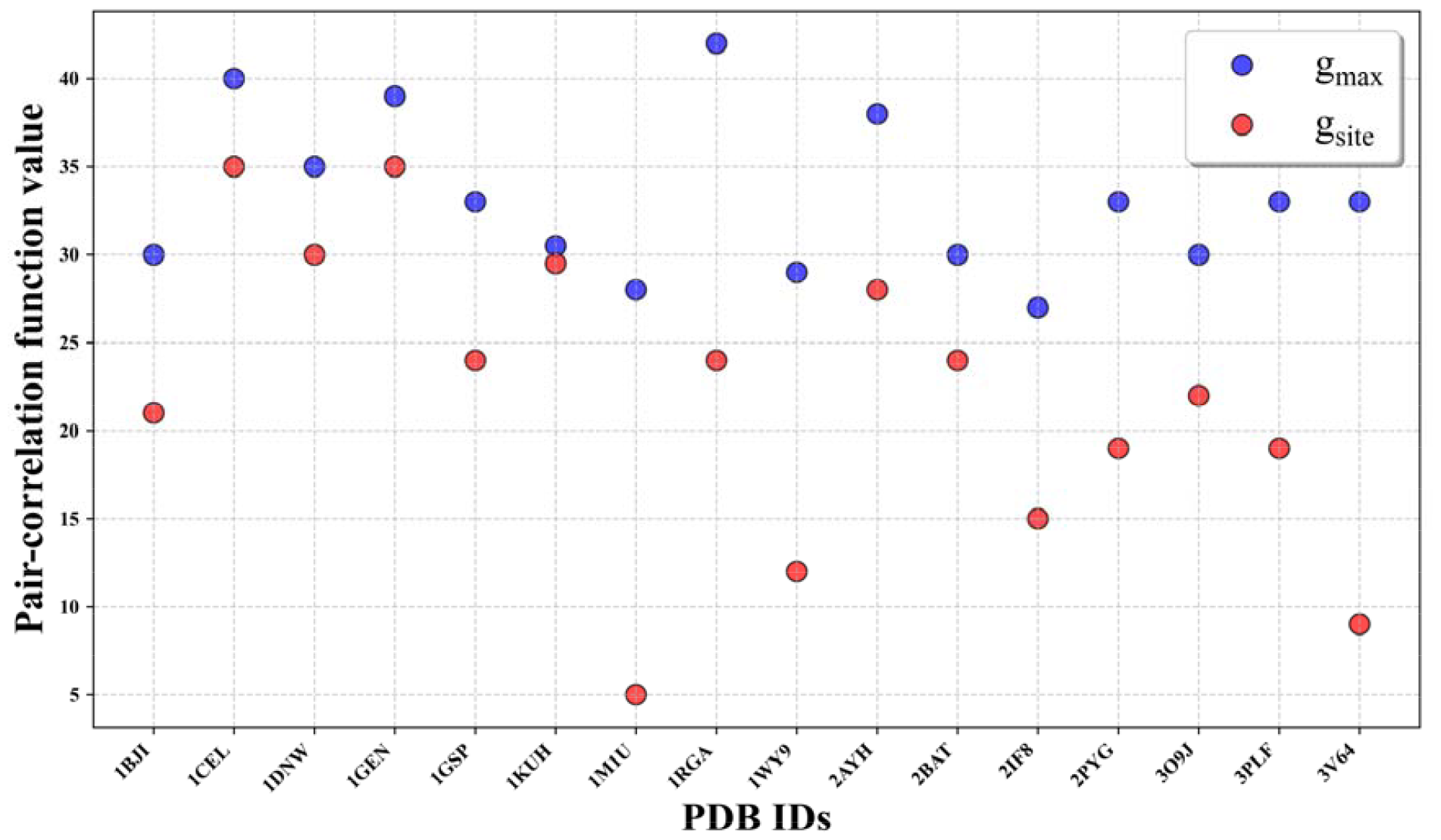
While we anticipate the presence of g_max_(**r**) in the binding site for a successful prediction, the algorithm does not dismiss the possibility of Ca^2+^ presence in the binding site, even in cases where the prediction fails. The figure compares the highest values of g(**r**) in protein binding sites, referred to as g_site_(**r**), with the maximum g(**r**) values, denoted as g_max_(r), for a protein from the unsuccessful cases.

### 3.3 Calcium Site Predictions: Assessment of the Binding Site Geometry

We employed the CheckMyMetal server to obtain insights into the overall modelling of calcium site environments ^78^. Our assessment focused on evaluating geometry and the completeness of geometry. Additionally, we examined the valence and vacancy of Ca^2+^ ions, as illustrated in Figure 6. In this context, “valence” represents the sum of individual bond valence values (v_i_) for each metal-ligand interaction within the coordination sphere, as shown in eq. (9). The term “nVECSUM” involves normalizing the magnitude of the vector sum of bond valence vectors by the overall valence. A bond valence vector is a vector with a magnitude equal to the bond valence and drawn from the central metal to its ligands. This calculation depends on individual bond valence vectors associated with atoms located within a 4 Å radius of the metal. In an ideal scenario characterized by perfect symmetry and consistent bond lengths, these individual bond valence vectors would effectively cancel each other, resulting in an nVECSUM value of 0 ^84^. Compared to their idealized values, the gRMSD is the root mean square deviation of observed ligand-metal-ligand angles where ligand means the atoms coordinating with the calcium ion. It is calculated individually for each metal-binding site, determining the geometry deviation value for each potential reference geometry as defined in reference 78. The determination of vacant coordination sites occurs after selecting the best-fitting geometry. The vacancy parameter is subsequently computed by expressing the percentage of these empty sites with the coordination number associated with the chosen geometry ^78^. We have compared the values of these five parameters for the successful versus failed cases.

**Figure 6:**
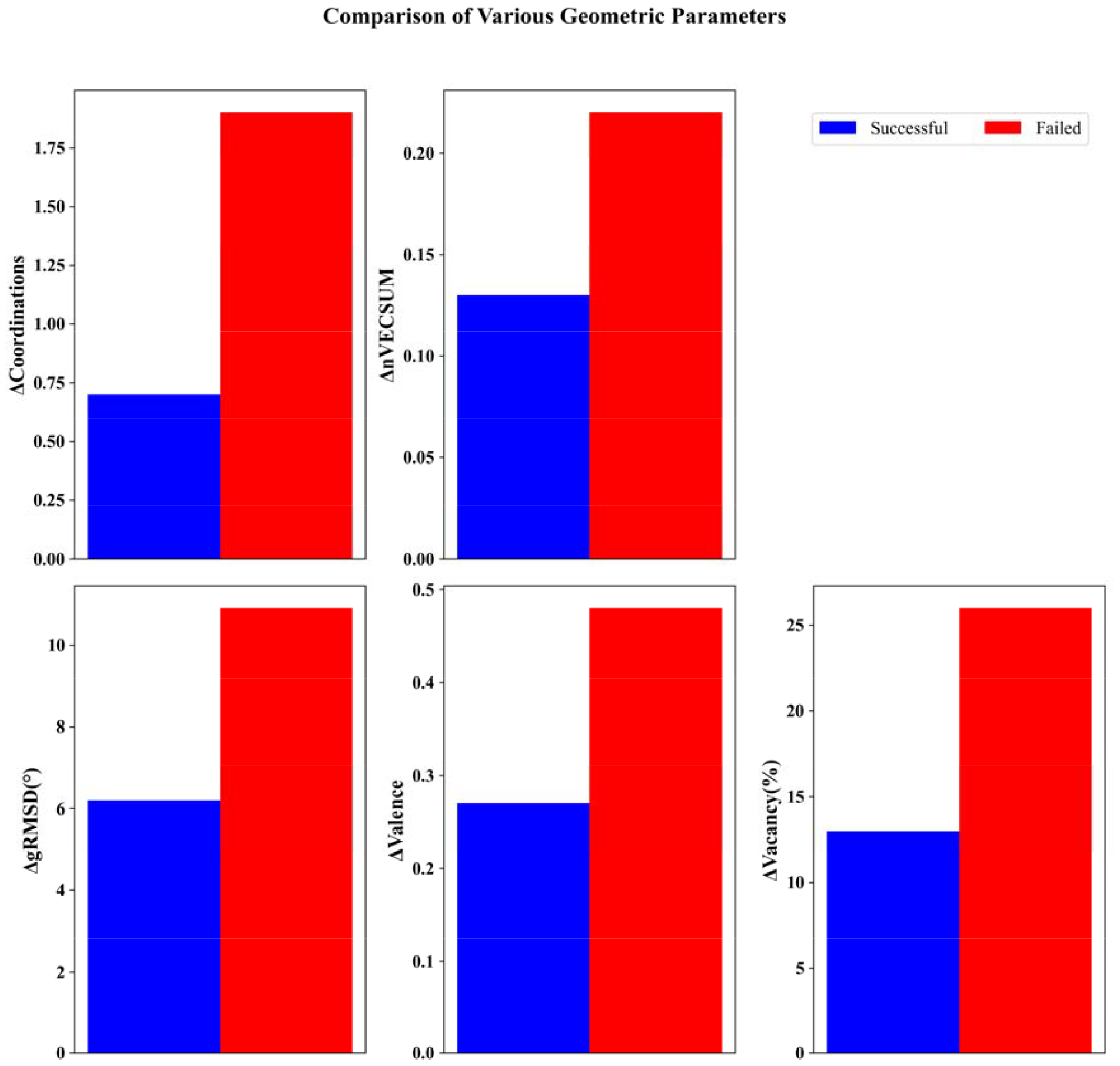
The comparison between the absolute mean difference in experimental and predicted geometric parameters for the successful and failed cases of calcium-binding proteins. Together, all these five parameters define the environment of a metal ion.

Figure 6 shows the average absolute difference and predicted and actual site parameters. The successful cases unambiguously show smaller deviations than the unsuccessful cases. For the coordination, the failed cases, on average, show more difference in coordination than the actual values compared to that for the successful cases. Differences in gRMSD indicate that in the failed cases, on average, geometry is more distorted. Overall, there is more average deviation in these five parameters for the failed cases; however, it is to be noted that the number of proteins in the successful cases is more than seven times that of the failed cases; hence, this analysis can give only a qualitative picture.

### 3.4 Analyzing Factors Affecting Prediction Accuracy in Algorithms: Structure Quality Assessment

We examined the quality of the protein’s structure in cases where the algorithm failed. This examination unveiled a common trend among the failed instances – a notable presence of low structural quality indicators, including parameters like R-factor, clashscore and Ramachandran outliers. Moreover, upon closer inspection, it became evident that several coordinating residues within these failed cases exhibited geometric irregularities or encountered challenges related to electron density interpretation. These findings suggest a strong correlation between structural quality and the accuracy of our algorithm. Seven of the sixteen failed cases have low resolution (≥ 2.0). Interestingly, in the five cases of the remaining nine cases from the group of failed predictions, the R-factors are notably high(≥0.20), indicating potential errors or inaccuracies in the structural models.

Researchers predominantly depend on “global” metrics such as resolution, R-factors, and model quality assessments for constructing models. Rigorous validation procedures rooted in global measures ensure the overall correctness of structures. However, these global measures are not well suited to enable users of crystallographic models to assess the dependability of “local” features in specific regions of interest. Consequently, even a crystal structure founded on high-quality diffraction data, meticulously built and refined, may exhibit less reliability in local areas than the rest of the model. Misinterpretation of this fact can lead users astray, particularly in mobile regions of the structure, including active sites, surface residues, and ligands.

After examining the global metrics (resolution and R-factors), we investigated site-specific factors. We analyzed the B-factors associated with calcium-coordinating oxygens atoms, narrowing the study to the remaining four cases. If the standard deviation of B-factors surpassed two times the standard deviation (σ) from the mean (μ), we considered these atoms’ heightened dynamics or uncertain positioning. None of the B-factors of oxygens in proteins exceeded the established cutoff (μ ± 2σ). Subsequently, we examined the validation reports accompanying the PDB entries ^94^ to assess the quality of residues coordinating with the Ca^2+^. We found one instance, i.e. 2AYH, where the coordinating residues exhibited geometric concerns (Figure 7). Then, we examined Ca^2+^ assignment after assessing protein structure metrics and site-specific factors. We used the COOT software to analyze the electron densities associated with the Ca^2+^ within the binding sites ^76^. We found a specific case, 3PLF, where, upon examining the electron density and negative difference maps (2Fo – Fc and Fo – Fc at 2σ level), we observed that the calculated electron density is more than the observed electron density at the Ca^2+^ site. It indicates that Ca^2+^ is probably missing from that site (Figure 7), and previously assigned sites in these structures need to be corrected.

**Figure 7:**
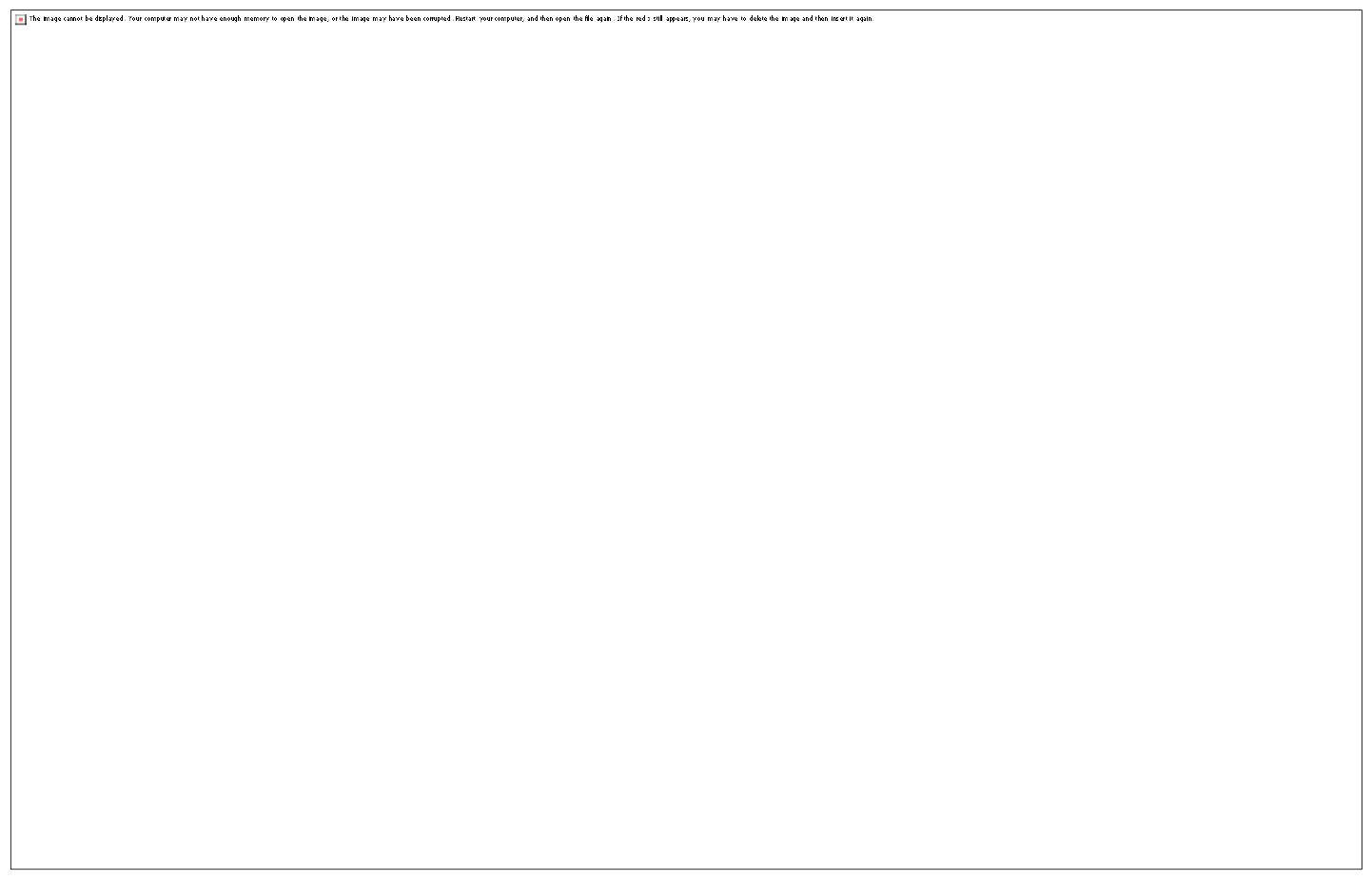
(a) Representation of a Ca^2+^ site (PDB ID: 2AYH) with a modelled angle of 126 degrees, surpassing the expected angle of 118 degrees. The anticipated bond angle is in blue. (b) Electron density surrounding the calcium-binding site in the MetRD peptide complexed with the c-Cbl TKB domain (PDB: 3PLF). We represent electron density in blue and the difference map in red.

Ultimately, our investigation extended to validating accurate metal identity assignments by applying the CBVS approach within the remaining two cases ^84^. Upon examination, we found an erroneous assignment of calcium at a site designated for sodium in these cases (Figure 8). In the case of 1CEL, coordination is with two GLUs and one water molecule. Notably, while the binding site is on the protein’s surface, our prediction placed it within a cavity. There are additional ASP and GLU in the second solvation shell of the g_max_(r) position. Interestingly, Altman’s work predicted a position within proximity of 1.29 Å but with a very high rank of 18, i.e. it was their 18^th^ best prediction. It is significant because the binding site is prominently on the protein structure’s surface. We can attribute this disparity between the actual location of the binding site and our prediction to differences in binding site characteristics and the complexity of the surrounding structural environment. Its outcome underscores the critical role of precise metal ion assignment in achieving reliable and accurate structural interpretations.

**Figure 8:**
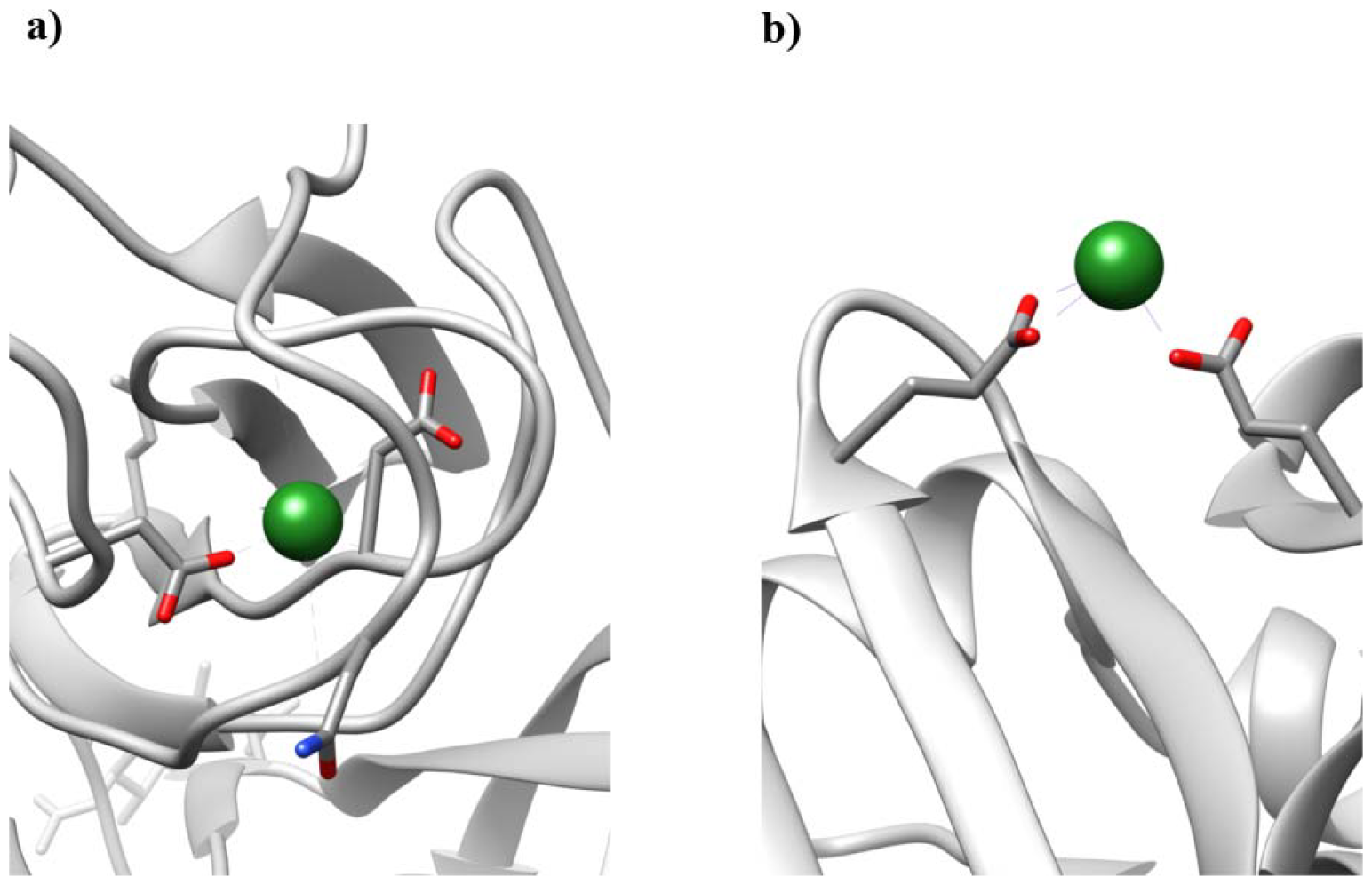
(a) Illustration of a case (PDB ID: 1BJI) and (b) case (PDB ID: 1CEL) showcasing instances where the Na^+^ site is identified as a Ca^2+^ site.

### 3.5 Calcium Binding Site Predictions: Insights from Structural Adjustments and 3D-RISM Calculations

The protein structures underwent unconstrained minimizations before 3D-RISM calculations. However, a pertinent consideration emerges regarding the potential impact of minimization on the binding sites or the overall geometry of the active site, potentially influencing subsequent 3D-RISM calculations. We subjected the structures to minimization under restraints on heavy atoms. The outcomes of this strategy highlight a noteworthy improvement in deviations; specifically, the mean deviation was notably reduced from 0.76 to 0.50. However, this was accompanied by a slight trade-off in the recall, resulting in 85%. This interplay between accuracy and deviation underscores the intricate dynamics that govern the prediction, with structural adjustments yielding commendable improvements in the deviation, albeit at a marginal cost to recall performance. Upon evaluating the structural quality of proteins in which predictions failed, we again observed similar issues as previously discussed in other structures. We conducted the subsequent 3D-RISM calculations following the minimization of the structure with positional restraints on heavy atoms.

### 3.6 Evaluating Prediction Performance on the Dataset-B

In the previous section, we identified issues related to the protein’s structural quality, like resolution R-factors and electron density in failed cases. Subsequently, we also examined whether the same problems were present in successful cases. Surprisingly, successful proteins exhibited similar structure-related issues. The success can be attributed to the robustness of our algorithm, which is likely to work even if structural irregularities are present. We filtered out all the problematic structures to get good models and structures to pinpoint the reasons for the algorithm’s failure, resulting in nine cases out of 134. We extended the assessment of prediction performance to the refined subset of previous proteins, i.e. Dataset-B. Remarkably, the prediction on this dataset showed a recall of 100%, indicating the ability to identify all binding sites correctly. Equally significant, the mean deviation metric was further reduced to 0.34, signifying enhanced prediction precision. We have examined the effect of individual filters on the success rate of our algorithm. The following table shows the impact of various structural filters on the protein structures in the dataset, demonstrating the reduction in the number of structures as each filter is applied.

According to Figure 9, our findings exhibited noticeable enhancements when applying a global filter, i.e. resolution and R factors. However, the transition to local filters introduced a non-linear relationship between results and filters. There is an issue of exceeding the B-factor in only one structure; using the third filter, i.e. the B-factor, results in negligible effective outcome changes. Subsequently, analysis involving validation reports identified problems in coordinating residues. Upon filtering out structures associated with these issues, recall decreased, but mean deviation improved concurrently. Continued refinement through filters focused on electron density around the calcium site and CVBS approaches led to further improvements in both recall and deviation.

**Figure 9:**
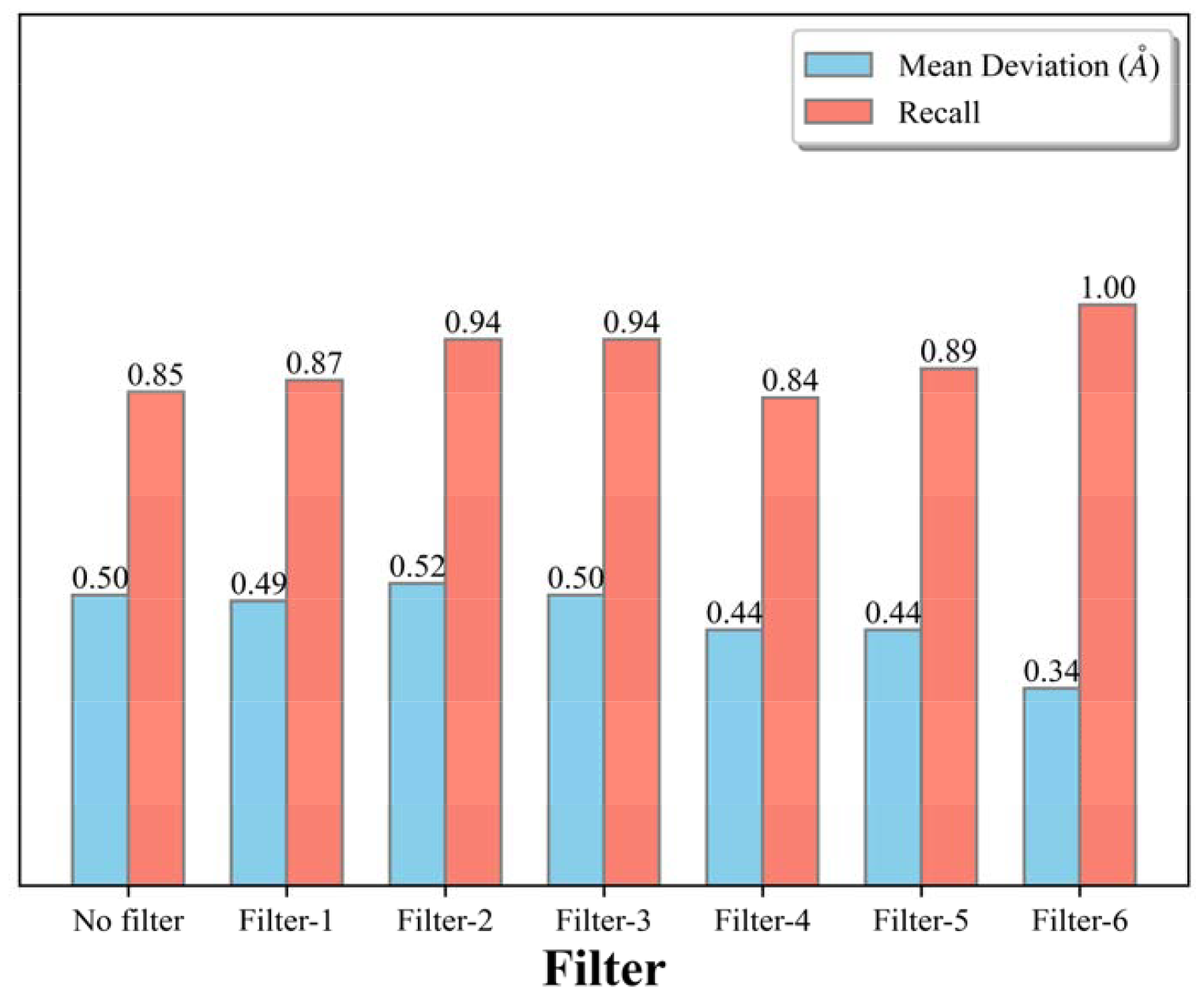
Comparison of Mean Deviation and Recall for Different Filters. The bar plot illustrates the mean deviation and recall values for six different applied to dataset-A. Each bar represents a specific filter of Table 1, with values labelled above.

### 3.7 Assessing Prediction Algorithm Performance on the Dataset-C

Then, we further evaluated our prediction algorithm on a high-quality protein dataset, which revealed a recall of 85 %, indicating the algorithm’s proficiency in identifying a substantial portion of actual binding sites. The mean deviation, quantifying the positional accuracy of the predictions, stood at an encouraging 0.40. Interestingly, a notable observation emerged even within the instances classified as prediction failures. Significant g(**r**) values were consistently detected at the intended binding sites, implying that the algorithm could capture the potential binding locations. Figure 10 illustrates the outcomes across three distinct datasets: A, B, and C. Two crucial metrics, mean deviation and recall, characterize each dataset. The filtered dataset stands out with the highest recall score of 1.00, emphasizing its proficiency in retrieving a significant proportion of relevant data.

**Figure 10:**
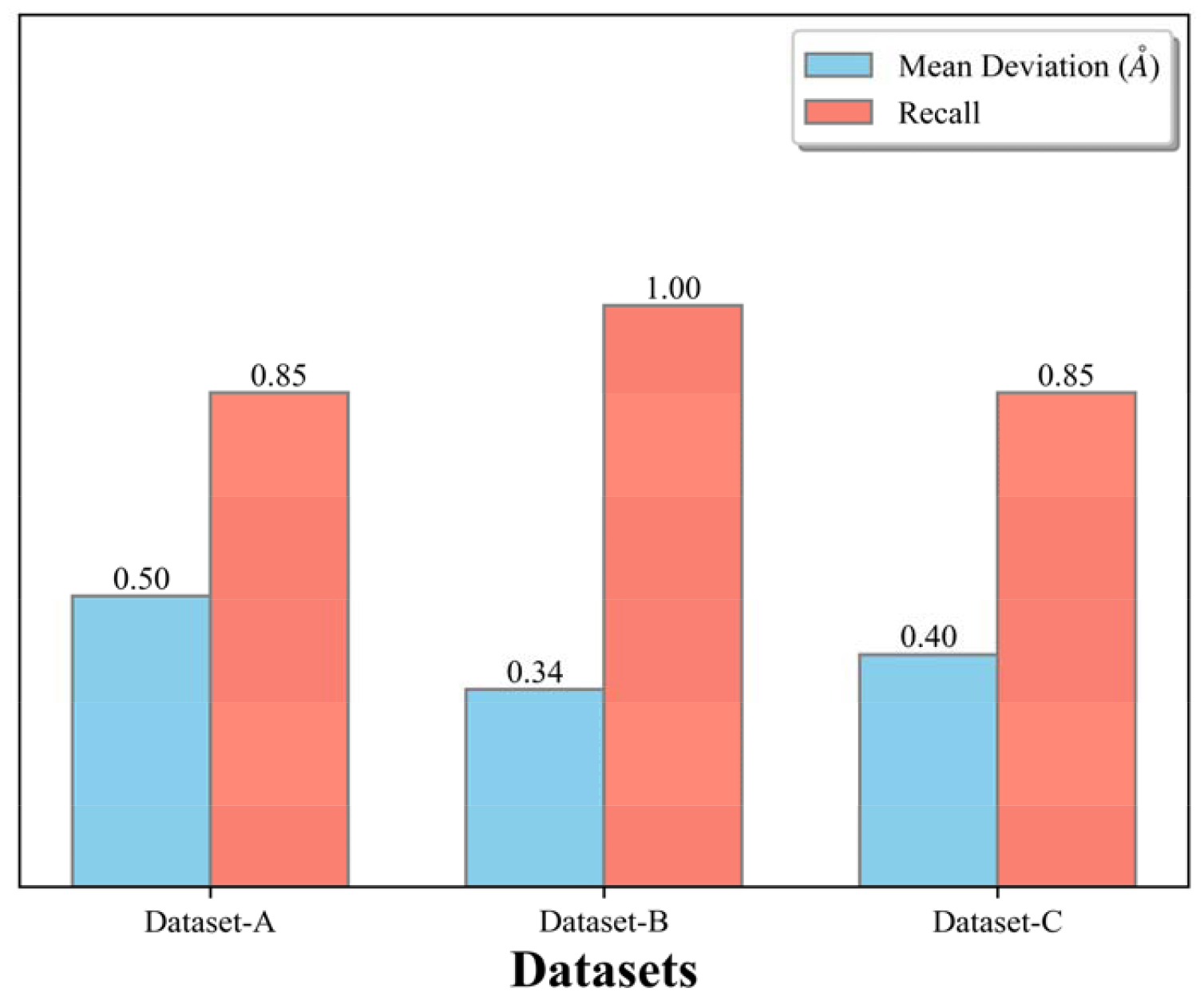
Comparison of mean deviation and recall across different datasets: The bar plot illustrates the variations in mean deviation (sky blue) and recall (salmon) for three datasets— A, B, and C.

## Discussion

### 4.1 Comparison with FEATURE

Using the FEATURE software, Altman and colleagues aimed to improve calcium-binding site prediction in protein structures ^42^. They trained the model on a dataset of known protein structures with verified calcium-binding sites, extracting features like local environment traits and structural motifs. The FEATURE-based method demonstrated inadequacy in predicting the sites when ligands other than calcium were present, as illustrated in the case of 1AG9. Consequently, we chose to exclude these structures from our datasets. When the calcium site exhibited unusual characteristics or lacked expected features, their approach failed to detect the site (e.g., 2TMV). Similarly, in our cases, our algorithm encounters difficulties in detecting sites with low coordination or a deficiency of negatively charged residues. Altman achieved an FRP of around 80% in his work. In our study, the FRP was slightly higher, reaching 84% for initial, 100% for filtered, and 73% for high-quality data. Our predictions show a somewhat increased FRP ratio compared to Altman’s results. Their recall is around 90%; in our study, the recall is reported as 85% when considering the initial and high-quality data. Notably, our recall reached 100% for the filtered data, indicating that our approach successfully identified all positive instances. Moreover, our precision surpasses theirs, as indicated by median deviations of approximately 0.30. This performance compares favourably to their reported figures of around 0.5 Å.

### 4.2 Water Network Analysis around the Ca^2+^ binding sites

One additional advantage of our methodology is that we can get a water network surrounding the Ca^2+^ ion. Figure 11 illustrates three representative proteins where water is present in the calcium-binding site. Typically, only oxygen atoms from water are present in these sites. Our analysis focused on comparing the positions of these oxygen atoms to understand their spatial arrangement and potential implications in calcium binding. Notably, local maxima in the hydrogen densities, i.e. g_H_(r), are predominantly situated near polar residues, highlighting the theory’s ability to discern and emphasize the influence of these residues on the hydrogen bonding patterns in the hydration process. In rare instances, hydrogen atoms of water are present in the X-ray structures in the PDB. Therefore, we showcased the distribution for g_H_(**r**) > 4 only, highlighting specific cases where hydrogen is present as part of water at the calcium-binding site.

**Figure 11:**
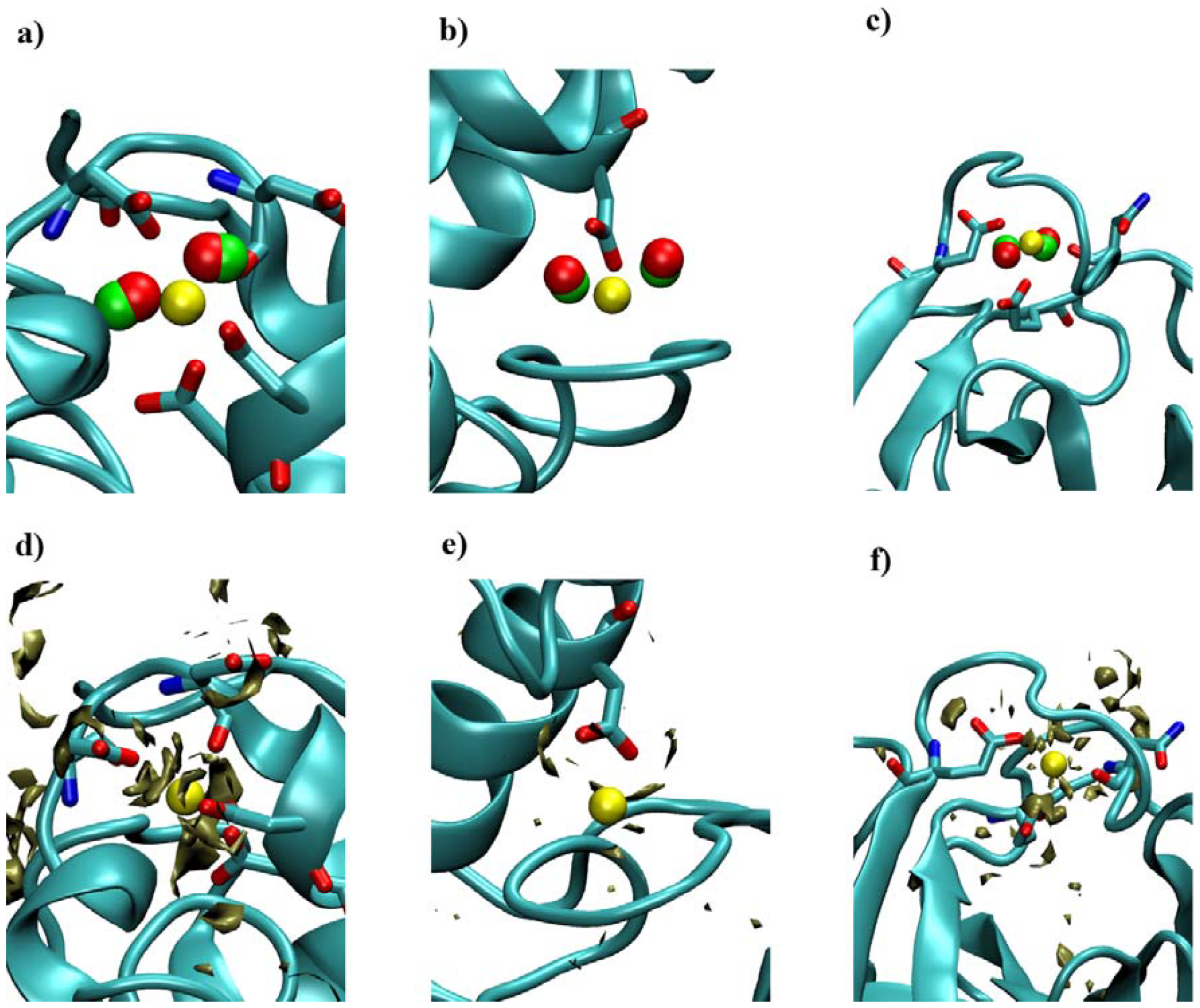
The upper panel of the figure depicts the crystal oxygen from water molecules (in red) and predicted oxygen positions (in green) surrounding the Ca^2+^ binding site in the proteins, identified by their respective PDB IDs: 1B9O, 1BP2, and 3LJJ. The calcium ions are yellow. Notably, these cases’ positions of oxygens and Ca^2+^ are concurrently determined. The lower illustrates the spatial distribution of hydrogen density surrounding Ca^2+^ binding sites of the same proteins. Notably, areas with an elevated concentration of hydrogen atoms, depicted in tan colour, correspond to regions around polar residues.

The binding sites of Ca^2+^ ions within proteins exhibit a remarkable irregularity, characterized by a wide array of coordination numbers. This intrinsic variability sets them apart from other metal ions like zinc and iron, which often adhere to more predictable coordination geometries ^26^. Consequently, conventional approaches relying on homology or geometry may falter when applied to these Ca^2+^-binding sites. In light of this complicated landscape, the proposed method takes a holistic perspective encompassing the ion and surrounding water molecules’ complex network. The analysis focuses on cases where the binding site prediction algorithm failed Ca^2+^ binding proteins. The investigation revealed some primary reasons for these failures. A notable advantage of the physics-based approach is its capability to compute the effects of protein mutations or modifications without explicit training on databases tailored for such alterations. Our protocol’s ability to produce results within hours highlights its efficiency compared to the large computational cost for MD-based methods ^95,96^. This significant speed enhancement comes from the semi-analytic nature of the RISM theory as opposed to the sampling of waters and ions in an explicit-solvent MD simulation. Moreover, the convergence of ion distribution in MD simulation, even for a single protein structure used in this work, can take substantial time.

While our current model exhibits strengths, there are promising avenues for refinement. Firstly, the accuracy of these calculations hinges on force fields. The standard force fields overestimate interactions involving highly charged ions like calcium. One avenue for improvement consists of adopting polarizable force fields, especially the charge scaling approach used to model the interaction between Ca^2+^ and proteins ^97,98^. It’s crucial to acknowledge that the 3D-RISM method we employed to solve the Ornstein-Zernike equation has its approximations and relies on an empirical closure relation. In light of a handful of unsuccessful predictions, it is crucial to emphasize that the binding sites continue to exhibit significant g(**r**) values. This observation highlights potential interaction sites, suggesting that the model captures essential aspects of the underlying molecular interactions despite occasional discrepancies. Lastly, using the placevent algorithm for ion predictions leads to underestimation ^68^. Indeed, it is well-established that some calcium sites can bind potassium, sodium, and magnesium ^99^. Furthermore, numerous metal-binding sites can bind to multiple metal ions ^100^. Hence, it becomes imperative to ensure specificity or discrimination between Ca^2+^ sites and other metal-binding sites to enhance the accuracy of predictions. Continually exploring these avenues holds the potential for refining and advancing our predictive model.

This work has aimed to develop an innovative and fast algorithm grounded in statistical mechanics principles for identifying Ca^2+^ binding sites in proteins known for such capabilities. The algorithm’s efficacy was evaluated on a diverse protein crystal structure dataset, demonstrating its robust power across sequence variations, structural motifs, and intricate binding site geometries. A distinctive feature is its explicit incorporation of three-dimensional distributions of water molecules surrounding proteins, based on the 3D-RISM/KH theory. The algorithm is a tool for accurately identifying Ca^2+^ binding sites, holding substantial potential. This advancement enhances our understanding of Ca^2+^-protein interactions, paving the way for innovative therapeutic strategies and catalyzing progress in the field.

## Acknowledgement

A.B. would like to thank Dr. Rakesh Srivastava for valuable discussions. This work is supported by a Department of Biotechnology (DBT) grant (BT/PR45865/BID/7/1016/2023) awarded to P.B. This work was also partially supported by grants from the DBT (BT/PR/40251/BITS/137/11/2021) awarded to the Centre for Computational Biology and Bioinformatics, Jawaharlal Nehru University and the MATRICS grant from SERB (MTR/2021/000365) awarded to P.B.

