## Supplementary material for "Prediction of Ca^2+^ binding site in proteins with a fast and accurate method based on statistical mechanics and analysis of crystal structures": SI

**Table 1**: This table lists datasets and associated PDB IDs used in our work, encompassing a broad spectrum of protein structures.

| **Datasets** | **PDB IDs** |
| --- | --- |
| Dataset-A | 1AI4 1ALC 1AQH 1AQM 1AYO 1B9O 1B9Z 1BF2 1BJI 1BK9 1BP2 1BUD 1CE5 1CEL 1CLV 1CLX 1CP7 1CVL 1DNW 1EGZ 1EL1 1ELT 1EX9 1EZR 1F8V 1FE5 1FNZ 1GCA 1GCG 1GEN 1GSP 1H30 1HML 1IAG 1IRB 1J0H 1J0M 1J5U 1J8E 1JC9 1JHN 1K12 1KE2 1KEC 1KUH 1KVW 1KVX 1M1U 1MHL 1NAE 1NBC 1NUF 1OIL 1OPO 1OZ6 1PNK 1PZY 1QFK 1QGE 1RGA 1SGT 1SMD 1SUS 1TGM 1U4G 1VB9 1VEM 1VSI 1WY9 1XD1 1XJO 1YU6 1Z3U 2AYH 2BAT 2BOH 2DDU 2DUO 2E81 2FIB 2H1U 2HET 2HIH 2IF8 2IPN 2JBK 2JHH 2MPR 2NN9 2NQX 2O1K 2PVS 2PYG 2QOG 2QPT 2QQM 2QQO 2STB 2TAA 2TGT 2VM9 2WG8 2ZKM 3A8R 3AIG 3BOD 3C7O 3CHJ 3D34 3EE6 3EST 3F6U 3G4E 3GBO 3GBP 3H4S 3LC3 3LHM 3LJJ 3M36  3O9J 3OHM 3PLF 3POB 3POS 3PRK 3PSR 3SH5 3V64 4BP2 4LIP 4PTP  4SGB 5CHY |
| Dataset-B | 1AQM 1B9O 1BP2 1CLV 1HML 1IAG 2IPN 2VM9 3LHM |
| Dataset-C | 1B9O 1GA6 1HJ8 1OD3 1QV1 1TKJ 1W0N 1W32 1YS1 2C1V 2FVY  2PWA 2WNP 2ZEX 3BEU 3LAG 3ZQX 3ZUC 4AAN 4ATE 4B9C 4B9F  4B9P 4I8H 4OP5 4OUS 4RU3 4XED 4Y0Y 5MFA 5U3A 5UQZ 6JK4 |

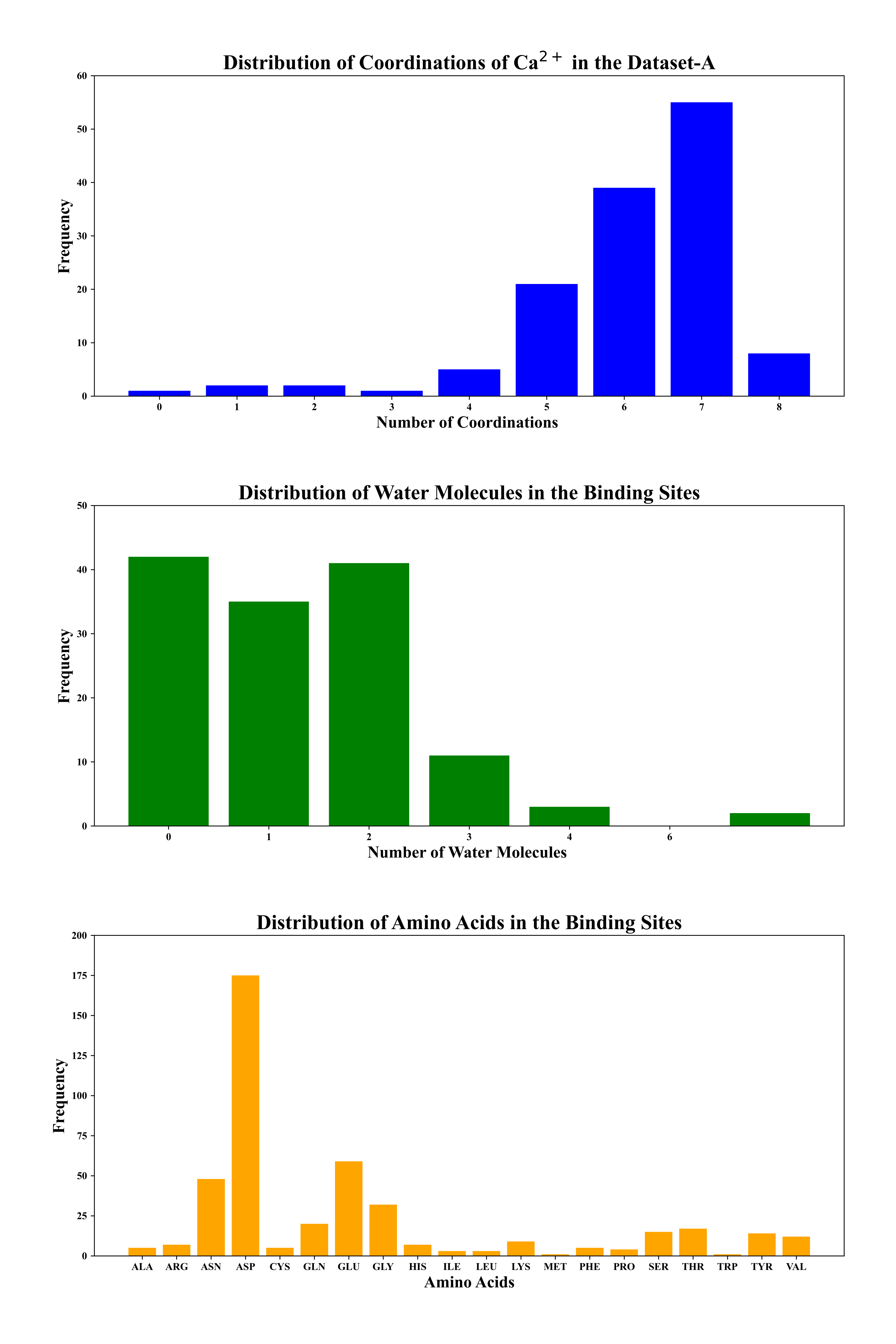

**Figure** **S1**: Illustrates the distribution of coordinations, water molecules, and amino acids in PDB entries in dataset-A. Each bar plot showcases the frequency of occurrences for different categories, providing insights into the structural diversity present in the dataset. Coordinations are depicted by the blue bars, water molecules by the green bars, and amino acids by the orange bars. The x-axis represents the respective categories, while the y-axis shows the frequency of each element.

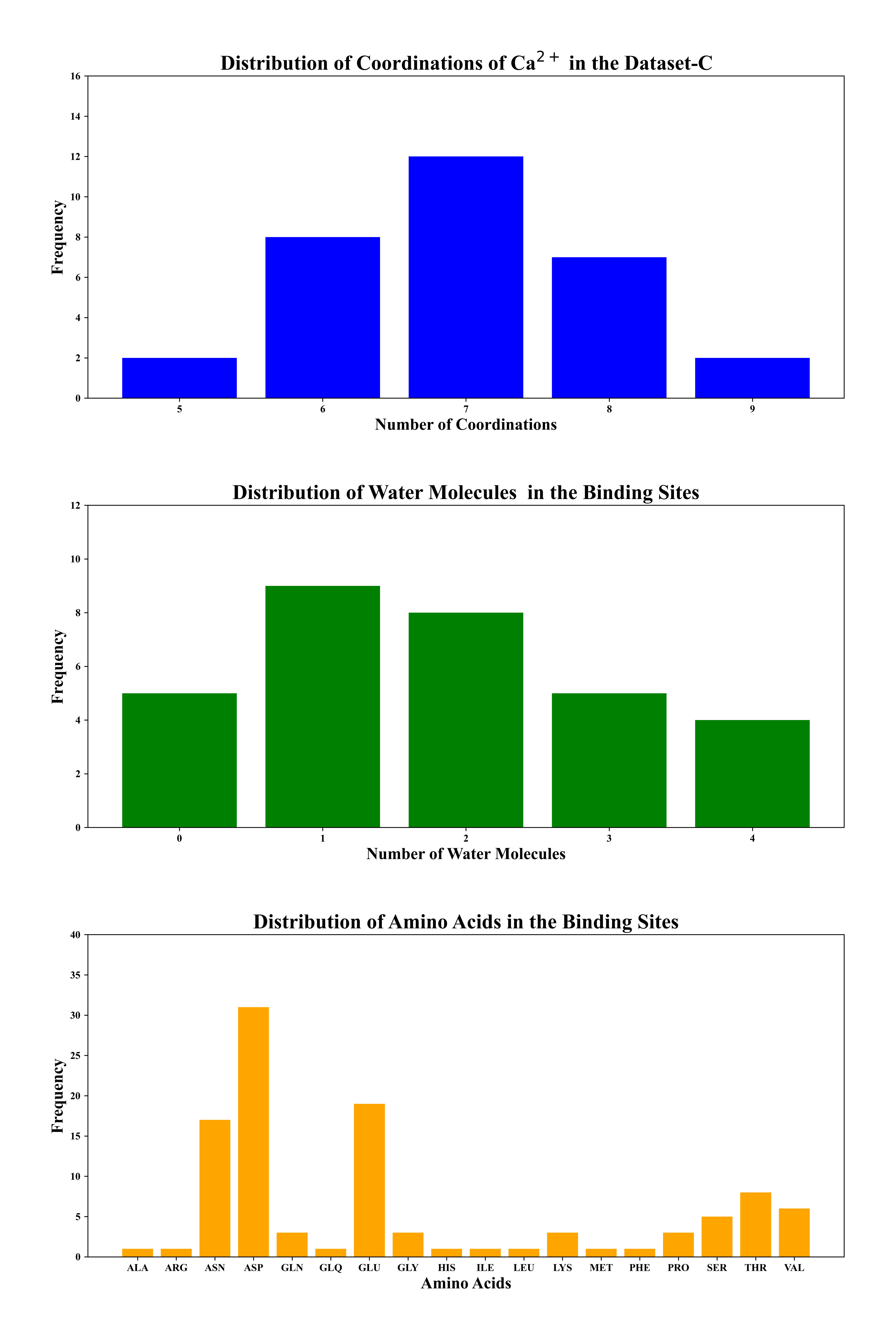

**Figure** **S2**: Illustrates the distribution of coordinations, water molecules, and amino acids in PDB entries in dataset-C. Each bar plot showcases the frequency of occurrences for different categories, providing insights into the structural diversity present in the dataset. Coordinations are depicted by the blue bars, water molecules by the green bars, and amino acids by the orange bars. The x-axis represents the respective categories, while the y-axis shows the frequency of each element.

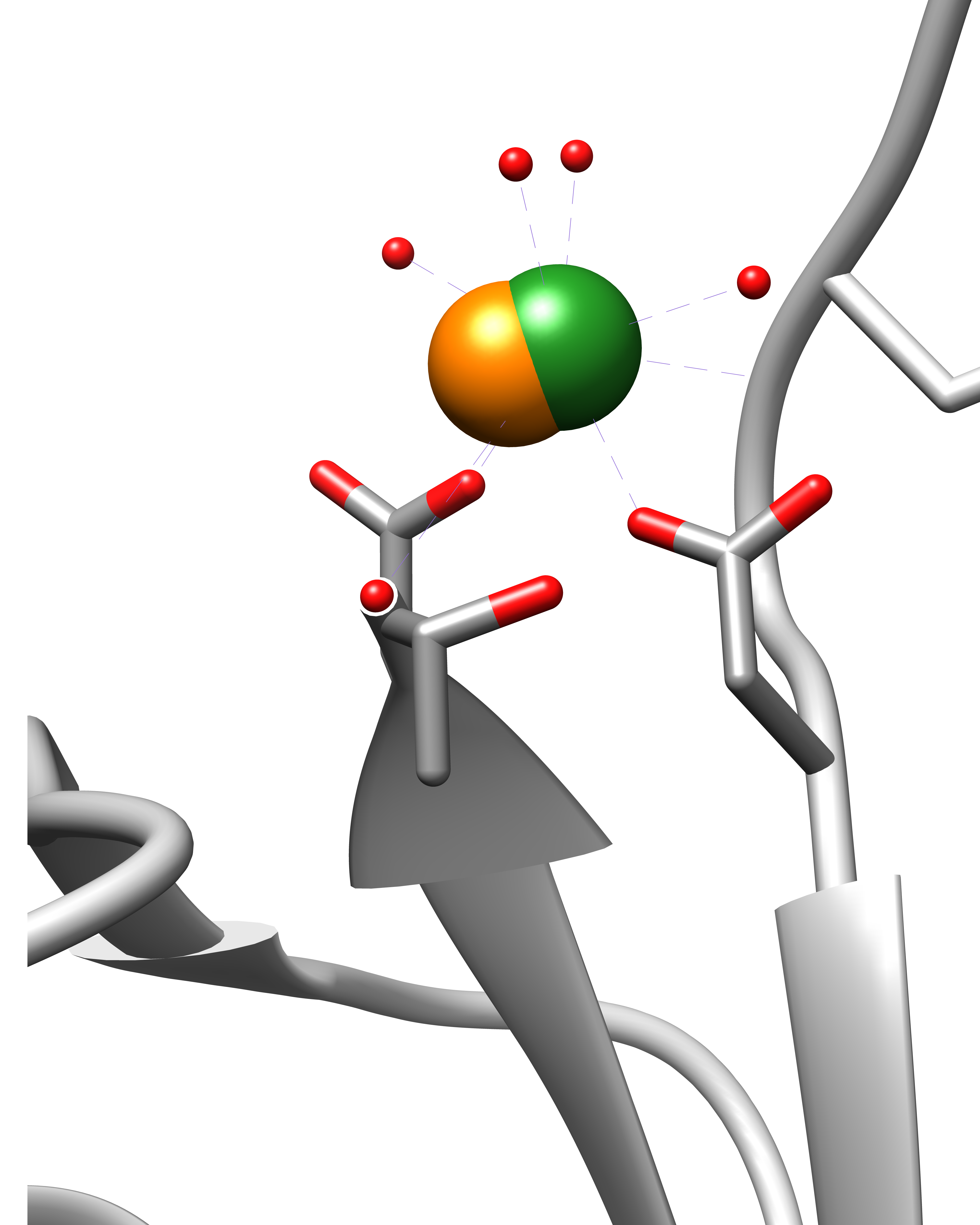

**Figure S3**: This figure presents an overview of 1J5U, where Ca^2+^ coordinates with four oxygen atoms. The crystal Ca^2+^ is depicted in green, while the predicted Ca^2+^ is represented in orange. Additionally, the crystal oxygen atoms of water molecules are red. This representation provides insights into the coordination and positioning of calcium ions with surrounding oxygen atoms from water in the binding site.
